# A framework for implementing metaheuristic algorithms using intercellular communication

**DOI:** 10.1101/2020.02.06.937979

**Authors:** Martín Gutiérrez, Yerko Ortiz, Javier Carrión

## Abstract

Metaheuristic procedures (MH) have been a trend driving Artificial Intelligence (AI) researchers for the past 50 years. A variety of tools and applications (not only in Computer Science) stem from these techniques. Also, MH frequently rely on evolution, a trademark process involved in cell colony growth. Generally, MH are used to approximate the solution to difficult problems but require a large amount of computational resources. Cell colonies harboring synthetic distributed circuits using intercell communication offer a direction for tackling this problem, as they process information in a massively parallel fashion. In this work, we propose a framework that maps MH elements to synthetic circuits in growing cell colonies. The framework relies on cell-cell communication mechanisms such as quorum sensing (QS) and bacterial conjugation. As a proof-of-concept, we also implemented the workflow associated to the framework, and tested the execution of two specific MH (Genetic Algorithms and Simulated Annealing) encoded as synthetic circuits on the `gro` simulator. Furthermore, we show an example of how our framework can be extended by implementing another kind of computational model: The Cellular Automaton. This work seeks to lay the foundations of mappings for implementing AI algorithms in a general manner using Synthetic Biology constructs in cell colonies.

## MAIN TEXT

Evolution is a key element involved in all microbiology processes. It is the process that drives organism adaptation to better survive and thrive in their surrounding environment. This process occurs at a genetic level, involving mainly genetic recombination and mutation. The genetic diversity produced by evolution is studied and used as inspiration in computational methods such as Evolutionary Algorithms (EAs)^1,2^. These algorithms are generally used for approximating solutions to optimization problems. Since evolution is a standard occurring process, it is natural to relate EAs to microbiology experiments, and more specifically, to Synthetic Biology constructs. This relationship has already been addressed by Directed Evolution^3,4^. However, the control level of Directed Evolution is not as specific as the one reached in EAs. Furthermore, several other computational methods can be translated to Synthetic Biology constructs that emulate their operation. Metaheuristic procedures (MH)^5–7^ are a larger class of procedures that contain EAs. Inspiration upon which these techniques are designed range from metallurgy processes^8–10^ through bird flock movement patterns^11–13^, and ant colony food foraging^14,15^. A general mapping, relating Synthetic Biology constructs to MH elements can be proposed such that any procedure of that class can be modeled as a synthetic circuit. This is due to MH sharing common elements and similarities that can be generalized.

This alternative paradigm for designing, implementing and executing MH is developed in the context of large-scale individual systems. The original population of solutions that take part in the execution is replaced by a set of individual entities, such as cells (in this work bacteria, specifically). Large-scale parallelism is a consequence of moving towards this new paradigm, but also the use of the procedure within a biological environment. This expands the scope of MH, establishing a wider array of possible implementations and problems to tackle. This association is logical, as these techniques work on a set of different elements (solutions) and apply changes on these elements to explore a search space and eventually reach a good solution in a reasonable amount of time according to specific constraints.

MH have been long studied and possess a defined structure^16^. One approach towards implementing AI using Synthetic Biology is shown in this article in the form of a framework that automates the mapping of MH elements to synthetic constructs. A proof of concept is implemented to show the automation of the process and generation of readily executable simulation files for the cell colony simulator `gro`^17,18^.

## RESULTS

One key aspect for using MH is to be able to represent all elements necessary for the execution of the procedure. Mainly, this involves the solution pool used in the execution, a fitness function to evaluate different solutions, and operations that carry out the exploration of new solutions. The application and design of these elements in a context of Synthetic Biology is not straightforward, as often they are dependent on the problem to solve. However, in this work, we propose a general mapping scheme to relate each of the elements which participate in an MH to a functional synthetic construct and make the association easier. The whole set of constructs is then organized and distributed over a pool of individuals (in this case, bacteria) to represent, and reproduce dynamics associated to the procedures. These constructs are designed from a general standpoint and seek to translate each of the involved components using transcriptional logic gates, intercellular communication mechanisms, and external elements such as environmental signals. It should be highlighted that the mapping presented here is a proposal and could be complemented and extended with other kinds of mechanisms, such as CRISPR^19,20^ systems, external conditions such as temperature, nutrient consumption, or specific spatial conditions. Also, it should be stressed that our proposal heavily relies on intercellular communication, since it offers a higher computation power and also distributes it among the colony cells. The main intercell communication processes used were bacterial conjugation^21–23^ and Quorum Sensing (QS)^24–27^.

The framework is composed by three parts:

1. A set of parameters that configure the execution of the instance of a MH. This set of parameters is always the same for the selected technique, despite having specific values to solve different problems. At this stage, the input parameters for the procedure are abstracted and generalized for multiple instances of the selected MH.
2. A mapping/translation language to relate specific elements of the MH technique to genetic circuits. This is the fundamental idea and value of the presented work, as it provides the blueprints for automating the design of MH in Synthetic Biology. How specific elements are ported to a genetic circuit will be discussed further in this section.
3. An interpreter that automates the translation of the specification of the algorithm into a skeleton of a `gro` source code, so the MH can readily be simulated and tested. The output design of this interpreter is generated based on 2) and configured based on 1). A depiction of the framework elements and their relationship is shown in Figure 1.

**Figure 1.**
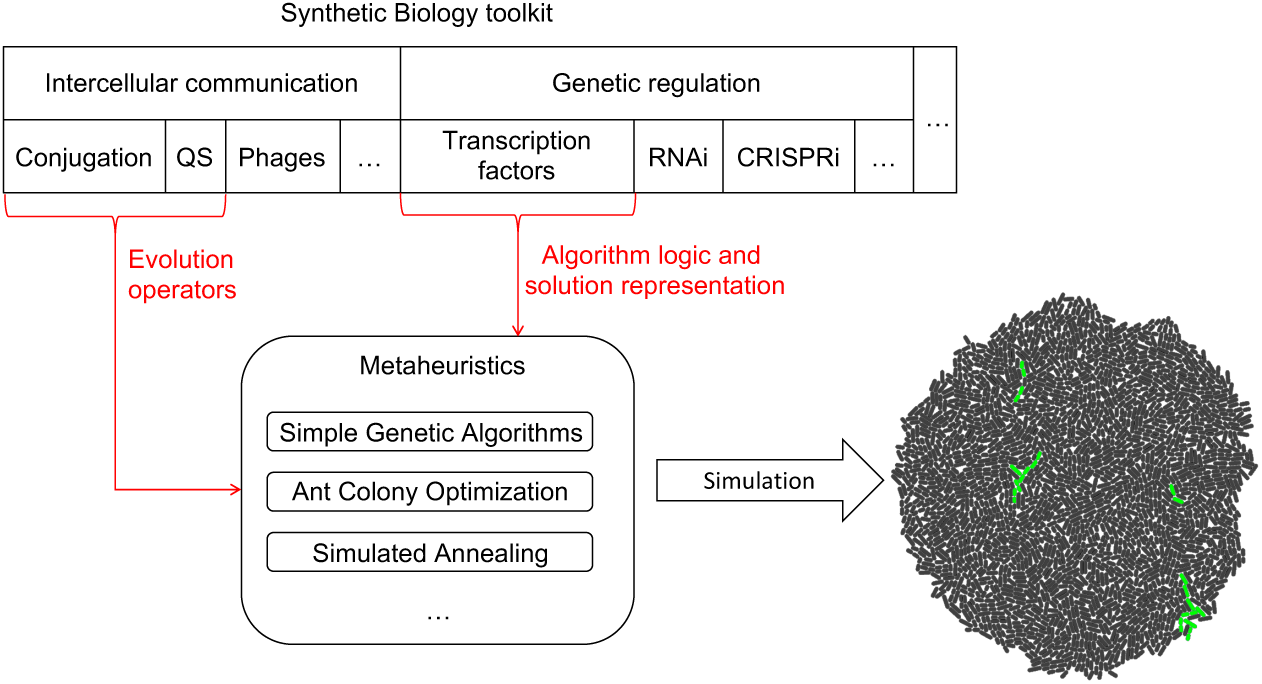
Framework elements. Intercellular communication processes were chosen as the tools to implement evolution operators in MH for cell colonies. Depending on what MH is implemented, each process could play a different role. MH logic and representation are encoded using transcriptional regulation. However, other possibilities such as the use of RNA or CRISPR mediated regulation remain open to be explored as new forms of logic and representation for MH. The selected tools are then mapped to the logic of the specific MH, generating a model. Once the model and mapping have occurred, a `gro` simulation file is outputted and run to analyze the behavior of the algorithm.

The aim of our framework was to generalize how MH are defined in terms of their parameters, establish base circuits which can be extended to generically model key players in MH procedures such as fitness functions, pool of solutions, or operations. We think this is a two-fold contribution as first, it eliminates the need for fully understanding all intricate mechanisms of the MH despite being able to use it, and second, automates its design thanks to the mapping that translates all of the elements into gene circuits (that are outputted in the form of a `gro` specification file, but also set a starting point in the design of gene circuit implementation related to the MH in the wet-lab).

We believe that each MH implemented for cell colonies following the proposed approach, and pursuing an optimization goal, represents a specialized form of Directed Evolution. It establishes further definition and control from an algorithmic standpoint, because the general algorithmic logic and evolution steps are explicitly specified. Furthermore, this continuous evolution is constantly being evaluated in MH by means of a fitness function. The variability for expressing and implementing this function within the context of our framework offers improved flexibility, expressiveness and specificity in the expected solutions with respect to the original definition of Directed Evolution.

Circuit design is done sequentially over the framework on the basis of fundamental part integration and the idea that all of the components of the MH procedure can be expressed in terms of these parts. Such parts merge into a more complex circuit that evaluates the inputs and outputs a function of these inputs. The circuits implement different elements of an MH, such as fitness function, solution representation, or evolution operators. These circuits will be presented and described after reviewing the basics on MH.

### Metaheuristics (MH)

MH are probabilistic techniques that take inspiration on certain observed general phenomena. The dynamics of the observed phenomena are then simplified and expressed in procedures that use input parameters for configuring and following the execution sequence. MH are mostly used for approximating solutions to difficult optimization problems. The simplification of the phenomenon and general approach of algorithm execution is a heuristic^28^. The heuristic is a function that seeks to guide the execution by estimating the reward that would be associated to carrying out a given step or strategy.

Inspiration for MH can originate from the most varied situations. Early instances of these techniques are Evolutionary Strategies^29^, Genetic Algorithms^30,31^ and Genetic Programming^32^. These procedures are all based on the phenomenon of natural selection. Evolving solutions, evaluating them and selecting for the best ones is the main heuristic driving these techniques. Since the evolution of the solution is guided by the heuristic, and the techniques are probabilistic, there is no certainty of convergence or eventually reaching the global optimum (and therefore are tagged as approximation procedures). However, the current best solution will tend towards an optimum (either local or global). Solutions are explored over a large landscape, called the search space. Since exploring the whole search space is not possible in most cases, strategies to partially visit the search space and evaluate solutions – to find the best possible one – are implemented. In sum, the procedure will actually be capable of improving the solution more by further evolving it, and in the long run (possibly infinite), will find the best possible solution. This same scenario can be modelled for other inspiration sources such as metallurgy processes, ant colonies or bird flock movements to name a few.

In the next subsections, the two MH (Simulated Annealing and Simple Genetic Algorithms) that will specifically be implemented and tested through our framework will be described.

### Simulated Annealing (SA)

SA^8–10^ is a MH that uses a controlled annealing process as inspiration for searching for optimal solutions. The goal of this algorithm is to best approximate a global optimum of a function. The definition of SA relies mainly on a temperature cooling function: the real annealing process requires the input metals to first be heated, merged, and later slowly cooled down. The goal is to achieve larger stable crystallization. Hence, the size and stability of the crystals depends on how the metal mixture is cooled down.

Being mainly an optimization technique, SA seeks to improve a solution according to specific problem conditions. The procedure works iteratively: first, a random solution is chosen and stored as the best one that has been found. Then, at each step, a random solution in the neighborhood of the previously chosen one is selected and evaluated. This new solution replaces the best one found with a given probability, and dependent on the temperature function and the fitness value of each of the involved solutions. This means that even a worse quality solution could replace the current best solution with a certain probability. The temperature function represents the probability of accepting any solution as a better one while the search space is explored. This function decreases as the algorithm execution progresses, entailing a gradually more localized search. The spirit of exploring potentially worse quality solutions is to reach other better solutions through them, and to not stay trapped at a local optimum.

The procedure of SA is shown in Figure 2.

**Figure 2.**
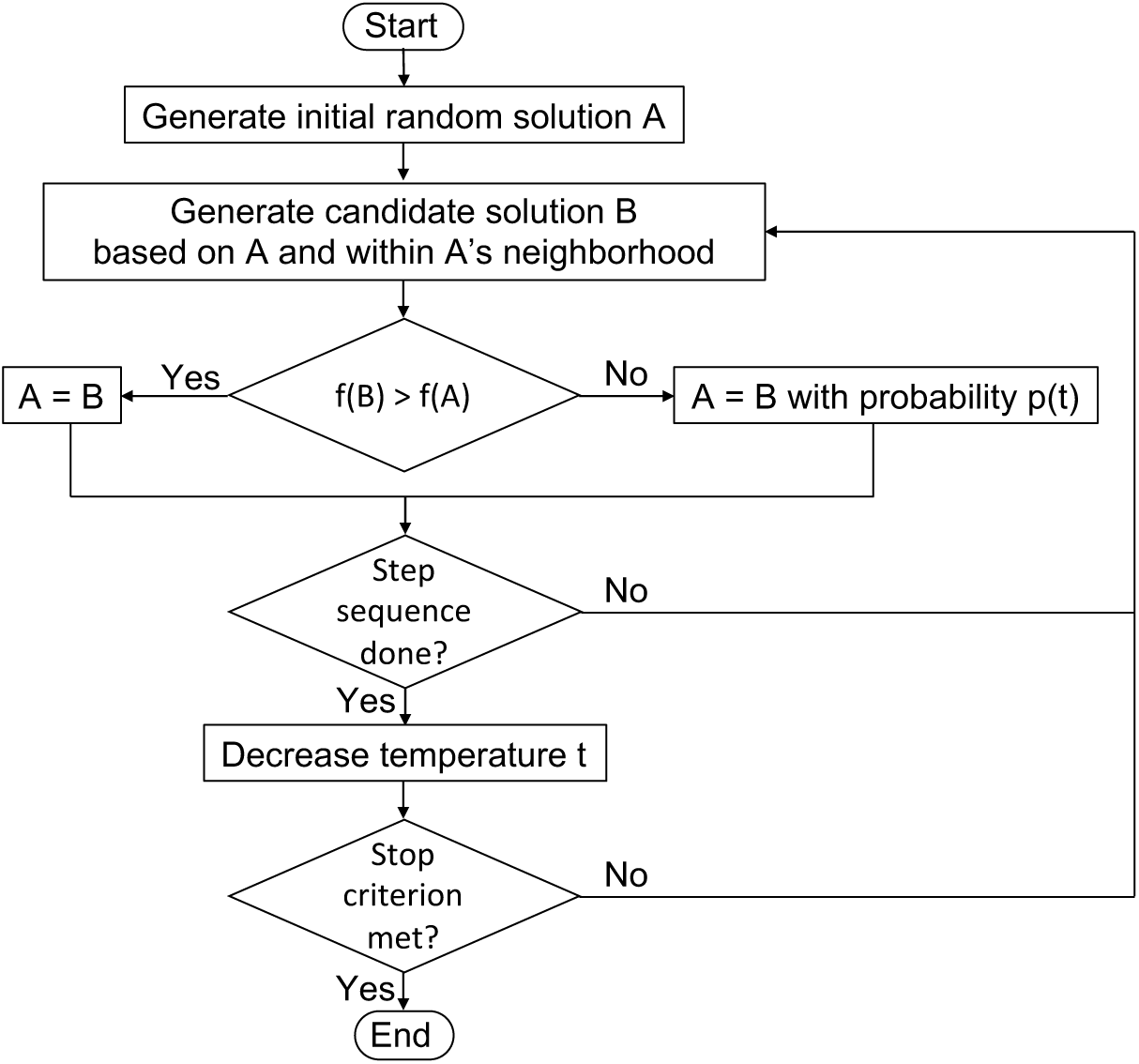
SA procedure flowchart. The candidate solution is first randomly generated and then evolved into a different one. Then, the new solution is tested for its fitness and replaces the previous solution probabilistically. The probability depends on a temperature variable, describing instability when the value is high, and translating into a higher chance of accepting worse solutions to explore different regions of the search space. The temperature variable is inspired on the real annealing process: it decreases gradually as the algorithm progresses in its execution. Examples of stopping criteria are number of iterations or fitness value reached.

### Simple genetic algorithm (SGA)

SGA^30,31^ finds its origins in evolution: it is an iterative MH that evaluates and evolves a pool of solutions in search of the fittest one. It is strongly based on evolution defined by Charles Darwin^33^. The key to assessing each solution lies in the definition of a fitness function, that returns a score based on the features of the individual being evaluated. Two kinds of operations are used for evolving: crossover and mutation. Each of these operations represents, respectively, local search (exploitation) and global search (exploration). Crossover is applied on a group of solutions it receives as input and returns a new group of solutions that are a combination of the former solutions. This operation seeks to model sexual reproduction. On the other hand, mutation is applied on a single solution and is defined as an arbitrary change in one or more of the traits of the solution. The goal of both operations is to obtain new solutions and preserve strong features of the explored ones in the case of crossover, or in the case of mutation, to increase variety of the solution pool. The search for a fit solution continues until the algorithm reaches a termination criterion. The basic operation of the algorithm is as follows: first, a pool of random solutions is created. Then, a subset of solutions is selected for reproduction (crossover) and generates new solutions with a given probability. Solutions in the pool then undergo mutation with a given probability. Finally, all of the solutions in the pool are evaluated using the fitness function, and a new initial pool of solutions is created for the next iteration. At this stage, termination criteria are also evaluated to check if the algorithm should stop. The procedure is depicted in Figure 3.

**Figure 3.**
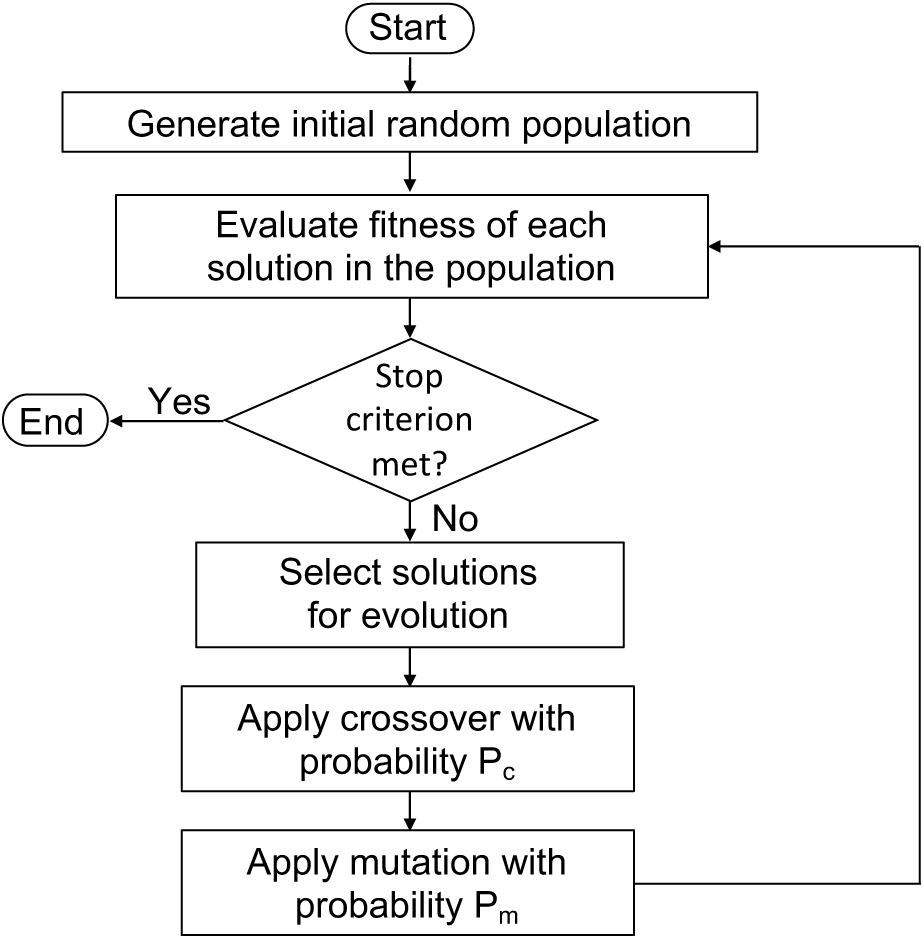
SGA procedure flowchart. SGA operates with solution populations, therefore, using cell colonies aligns naturally with its definition. Evolution is composed by three operations: selection, crossover and mutation. Furthermore, the solutions are evaluated at each iteration through a fitness function. All operations are applied until a stop criterion is reached. Typically, the stop criterion is met when a given level of fitness is achieved or when a certain number of generations (iterations) have been completed.

### Synthetic circuit designs for MH simulation

Heuristics are embedded in MH through a fitness function that evaluates and guides the search for best solutions within a search space. As a base assumption in this context, we will link the individual solution to the information inside a single cell. The presence or absence of a set of proteins of interest will act as the specific solution instance. Therefore, depending on which proteins are present or absent, each cell represents an individual and independent solution. Both evaluation and evolution dynamics will be implemented by taking advantage of cell capabilities. Bacterial conjugation will be used as the main backbone for evolution operations both in SA (solution mutation) and in SGA (crossover operation). Since solutions are represented as a set of proteins within a cell, perturbations of the set occur upon the arrival of a plasmid containing new proteins of interest into the cell. Fitness evaluation is organized in a synthetic circuit that senses the presence or absence of the proteins and performs a certain action (GFP expression in this case) when fitness is optimal.

A summary of the complete proposed mapping for both SA and SGA is depicted in Figure 4.

**Figure 4.**
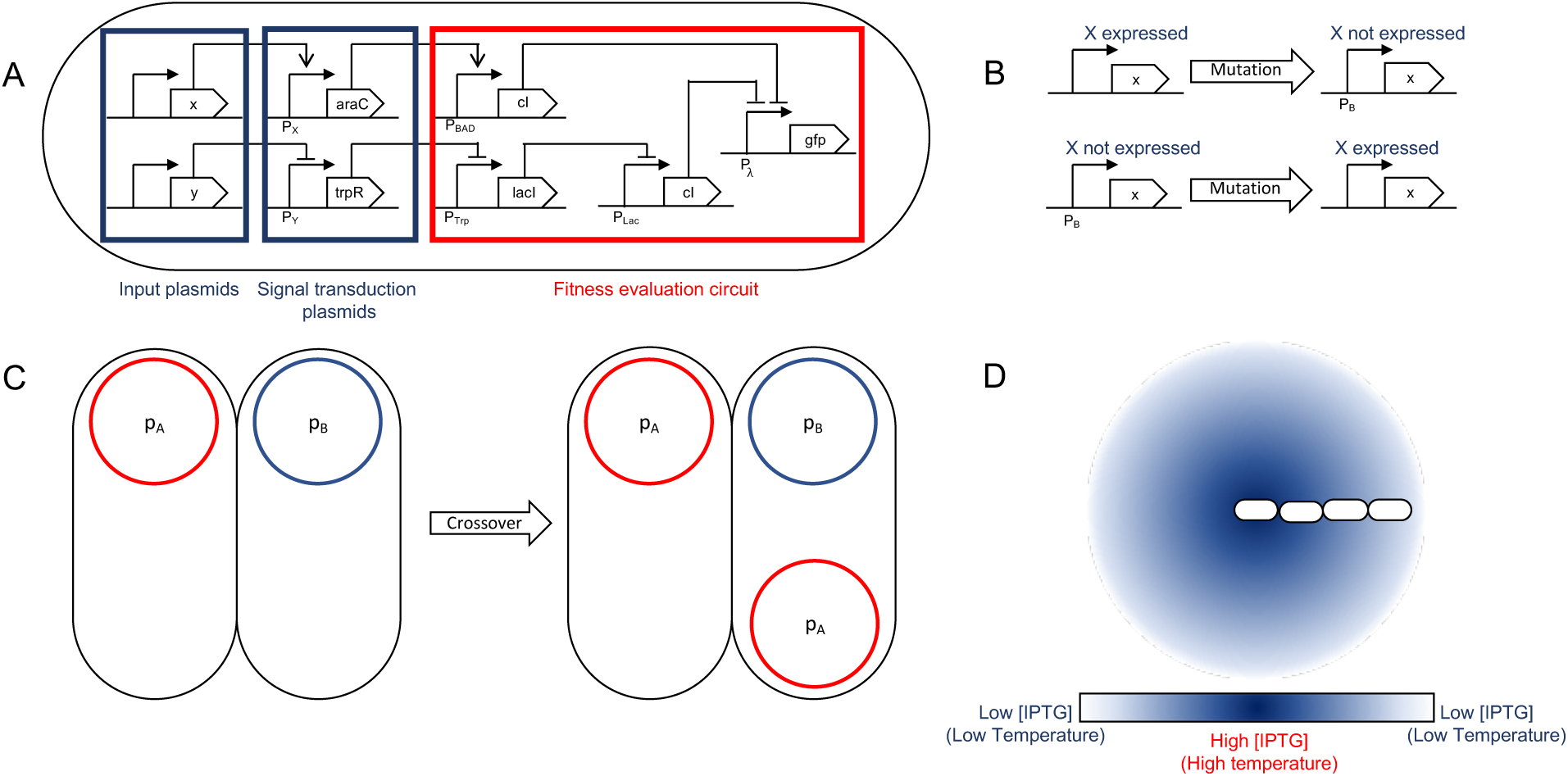
General logic mapping for SA and SGA. **A**. General gene network mapping for the MH framework. A three-tier design is proposed: the first tier are conjugative plasmids holding input proteins that are used by the fitness function to assess the quality of the solution. A second tier is a transition one in which the input signals are transduced into standard proteins for their evaluation. Finally, tier three is the evaluation circuit in which the input signals are checked against their respective set (if they need to be present, absent, or it is indifferent if they are present), and if said evaluation is successful, triggers an action. In this case, it is GFP expression, but that action may be replaced by any other. **B**. The mutation operation for SGA acts on the expression of a specific protein in the design, changing the solution to evaluate. The mutation rate parameter for SGA maps directly to the mutation rate configured in the simulation. This operation accounts for global search in terms of the solution exploration. **C**. Crossover is a recombination operation that we mapped to bacterial conjugation. Part of a foreign solution is integrated to the current one. In our mapping, we individualized a single protein to be held by a unique plasmid, therefore mobilizing a single protein between solutions for recombination. Conjugation rate is the parameter that accounts for the SGA crossover rate parameter. **D**. SA is largely based on a temperature decrease function: we use environmental signals (such as aTc or IPTG) as its representation. The temperature is associated to the signal concentration at a given location. The decrease is achieved by the shoving mechanical effects of the cell colony: the center of the colony experiences a greater temperature that the outer sections.

### Other evolution models

#### Cellular automaton (CA)

A CA^34–36^ is a n-dimensional grid structure where each of its cells has a state. These states are defined according to a set of given rules dependent on the states of its neighboring cells and the current cell itself. The rules describe interactions in the grid that alter the states and can describe spatio-temporal patterns in terms of the cell states. Based on the definition of neighborhood and specific rules involved in the design of the CA, it exhibits specific behaviors such as cyclic configurations or sequential movements. The most well-known early instance of this model is Conway’s Game of Life^37^. It is a binary CA in which grid cells die whenever their (Moore) neighborhood is overcrowded (more than 3 live neighbor cells) or undercrowded (less than 2 live cells), see Figure 5. Live grid cells stay alive when they neighborhood has 2 or 3 live neighbor grid cells. Finally, dead grid cells become live whenever they have exactly 3 live neighbors. This instance of CA will be the one inspiring our implementation. Although CA are not MH, the evolving nature of the model led us to test our framework logic, based on intercell communication, to implement CA.

**Figure 5.**
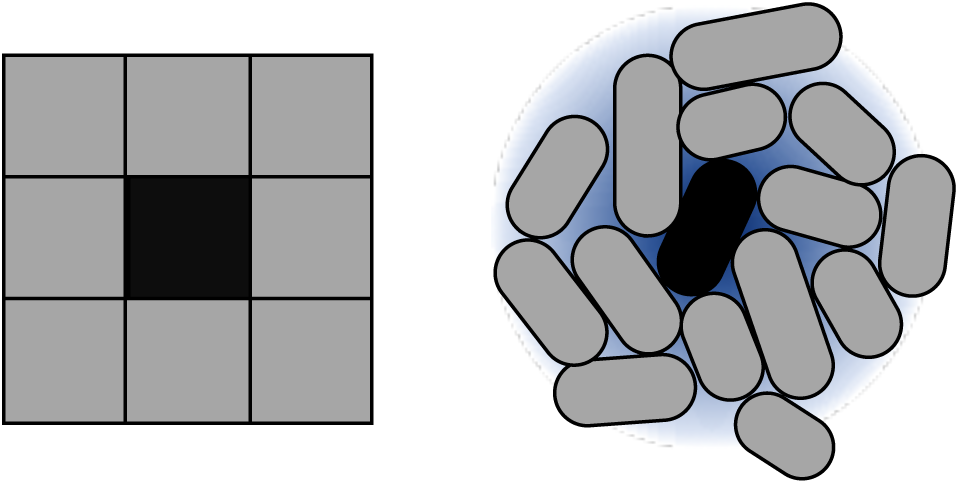
Approximation of Moore neighborhood using cells. CA execution is strongly dependent on the concept of neighborhood. 2D CA typically work on a Moore neighborhood. To reproduce this idea in the context of cell colonies, autoinducer sensing is used. The size of the neighborhood is dependent directly on the reach of the autoinducers. In terms of our model, this is represented through diffusion and degradation values of the intercellular signal.

### Implementation examples

To illustrate the capabilities and flexibility of the framework, we first present the phases involved in the parameterization, design and construction of the model along with the whole execution process associated with our implementation of the framework. Finally, simulation examples in the `gro` simulator are shown. Data for executions carried out for this work can be found in the Supplementary Information document.

#### Parameter collection for model generation

The first phase uses user-defined parameters to guide the shaping and automated generation of the model skeleton. Specifically, two of the three techniques mentioned above (SGA and SA) are examples that can be promptly executed in cell colony simulators using our implementation of framework. The third model, CA, is shown as an example of how it would be possible to extend the functionalities and models proposed by our framework through relating to the underlying models.

The fitness function and constraints associated to each of the MH (SGA and SA) are encoded through references to proteins and their interactions: since in `gro`, proteins are the unit for directing cell behavior, in our framework, they will mainly act as the base variables. The number of proteins used in the system is entered as a parameter, but also if each protein should be present, absent, or if it does not matter for describing a good solution to the problem. Concretely, the evaluation represents the fitness function. Also, the initial and final number of cells in the colony for the simulation are specified as additional parameters. It should also be noted that other proteins are used in the construction of the logic processes driving the algorithmic steps of each MH. Algorithm-related circuit construction process is automatically done in the next step.

These are all general parameters that are useful for specifying both SGA and SA. However, some specific parameters must also be collected in the case of each algorithm. For SGA, both mutation and crossover ratios must be provided. Mutation is implemented as an arbitrary change in the state of a protein within a cell, while crossover is simulated as a bacterial conjugation event. In the case of SA, the basic additional parameter is the temperature decrease ratio. In terms of `gro` simulations, this ratio is translated to the diffusion factor of an environmental signal (such as aTc), since the temperature can be associated to the signal concentration.

For implementing CA, they key parameter is setting intercell signaling using appropriate diffusion and degradation ratios. These parameters configure the distance from the signaling cell to its furthest neighbor. The goal of this configuration process is to emulate the Moore neighborhood in 2D. Within these settings, rules are encoded based on the concentration of signal sensed by cells. An example is shown in Figure 5, and a summary of the parameters involved in the framework (and for CA modeling) is compiled in Table 1.

**Table 1.** Fundamental parameters for modeling MH in cell colonies. Different parameters may be represented by the same biological equivalent depending on the MH to be modeled (conjugation rate, or signals for instance). All parameters marked for each MH need to be set for the execution of said simulation. This is crucial, because they control all aspects of the MH and require the specification in terms of their mapped biological equivalent.

| Algorithm parameter | Description | Algorithms | Biological interpretation |
| --- | --- | --- | --- |
| *Number of proteins* | Integer that specifies the number of proteins of interest for each solution | SA, SGA | Number of plasmids expressing proteins of interest |
| *Protein presence* | Contribution of each protein to the fitness function (if it should be present or absent) | SA, SGA | Expected expression state of a protein (ON/OFF) |
| *Initial cell count* | Size of initial solution pool | SA, SGA | Initial cell colony count |
| *Final cell count* | Size of final solution pool (stop criterion) | SA, SGA | Final cell colony count |
| *Mutation rate* | Determines mutation frequency | SGA | Promoter mutation frequency |
| *Crossover rate* | Determines recombination rate | SGA | Conjugation rate |
| *Temperature decrease rate* | Establishes size and cooling rate of the temperature zone | SA | Diffusion and degradation rates of an environment signal |
| *Solution perturbation* | Defines how the solution is altered for exploration | SA | Conjugation rate |
| *Moore neighborhood size* | Establishes size of the neighborhood of a cell | CA | Diffusion and degradation rates of an intercellular signal |

### Translation into the base model for simulation

Once the basic parameters and elements for the MH have been chosen and put in place, a model is constructed. This model summarizes the operation rules according to the specific mapping of the different elements present in the chosen technique to simulation instructions and constructs. The models are predefined and are an extensible (although specific) representation of the algorithms. Under our framework, it is possible to capture the essence and approximate the dynamics of MH through genetic designs. Therefore, as it is possible to model a MH based on genetic circuits, it can also be simulated.

The core representation of these circuits lies in protein expression that each define the elements of the solution for all MH. Therefore, a candidate solution is linked to a set of proteins being expressed jointly within the cell (shown in Figure 4). Intercell communication methods such as QS, or bacterial conjugation serve the purpose of providing a backbone for implementing operations on the existing solutions and obtaining new solutions. Specifically, under our representation, a set of plasmids hold the solution elements and their mobility aids in the dissemination of these elements within the colony. Since intercell communication is programmable, specific behavior regarding these operations can represent different operations for each MH.

### Automated generation of gro skeleton simulation file

Finally, with the model in place and informed by the input parameters, a simulation file generator constructs a `gro` simulation skeleton file. The mapping to the `gro` file takes place using the abstractions present in the simulator such as proteins, plasmids, environmental signals, etc. This file can be directly run by the `gro` simulator, or it can be modified by the final user for more specific operation.

### SGA execution examples

SGA examples were designed and implemented for simulation. In all examples, each solution is represented as a set of proteins that can be either present or absent. Each of these proteins is under the control of a promoter as a single gene in an operon. In turn, each operon resides in a different conjugative plasmid.

Each solution is then evaluated through a fitness function. This function is encoded in an operon that checks for a subset of necessary proteins that should be present, a subset of detrimental proteins that should be absent, and a subset of proteins that have no effect on the fitness of the solution. The operons that implement this function are also encoded in a single plasmid. Cells that comply with the requirements of the fitness function are classified as optimal solutions. Optimal solutions are marked by expressing GFP, and all other bacteria are uncolored. This is done merely for simulation purposes, but GFP could be replaced with different processes such as cell death, growth rate configuration or intercell signaling, depending on the purpose of the evaluation.

Crossover operation is mapped to bacterial conjugation between cells. Conjugation rate is therefore associated to the crossover rate parameter of the original SGA. Mutation operation was modeled as promoter mutation leading to incorrect functioning of the circuit, and arbitrary change in protein expression. Selection is random, since arbitrary recombination occurs, and bacterial conjugation is a simulated as a stochastic process.

### SA execution examples

Our team implemented simulations to test SA. Like in SGA, each solution is represented as a set of proteins of interest, and the fitness function also evaluates a solution based on the present/absent proteins in a cell. Concretely, a set of conjugative plasmids each holding a protein of interest, account for the values needed for evaluation each solution. The choice of conjugative plasmids to hold each of the proteins was made to better relate to a neighborhood space among solutions. It should be stressed that this does not refer to a physical neighborhood, but to a logical one for the solution set. If a set of proteins is represented as a binary string, binary neighbors of said string can be reached through conjugation.

Fitness evaluation is measured with a function based on interest proteins. A subset of proteins must be present, another subset must be absent, and it is indifferent if a third subset of proteins is present or absent. Similar to SGA, we established a color code in which green glow marks cells containing an optimal solution. Uncolored cells describe a solution with few or none of the necessary proteins being expressed. It should be stressed that, unlike the original definition of the algorithm, we did not implement plasmid loss (a possible way of finding a different solution), but only relied on plasmid mobility and aggregation.

The last important element of SA, the temperature value, was related to an aTc global signal. Therefore, this value is linked to the concentration of aTc at a given location. Temperature decrease is simulated mechanically, as cells are pushed outwards and experience a lower aTc concentration (equivalent to a lower temperature).

### CA execution example

Finally, and as an extension to the proposed framework, we implemented a simulation in `gro` relying mainly on QS. This was necessary to map the idea of Moore neighborhood to a cell colony context (see Figure 5). To preserve a static neighborhood, we eliminated growth from the colony simulation. The implemented CA simulation was an adaptation of Conway’s Game of Life. The synthetic circuit to implement this logic is based on the idea of band detection^38,39^: overcrowding and under-crowding are conditions that induce grid cell death, while a mid-level crowding amount induces grid cell life. The equivalence between the original Game of Life model and the proposed simulation is that a grid cell is mapped to a single cell in a colony.

The color code for this simulation was to use RFP for live cells and uncolored cells for “dead” ones. Cell state is determined based on the concentration of AHL at the cell location: high and low concentrations induce the cell to the “death” state, while a mid-level concentration makes the cell glow red. There are also some green cells, which have the task of starting the system, since there is no initial amount of AHL in the environment. These cells are placed randomly in the colony and are controlled by an environmental signal (aTc) as a start switch.

A summary of the circuits implemented for the simulations is depicted in Figure 6.

**Figure 6.**
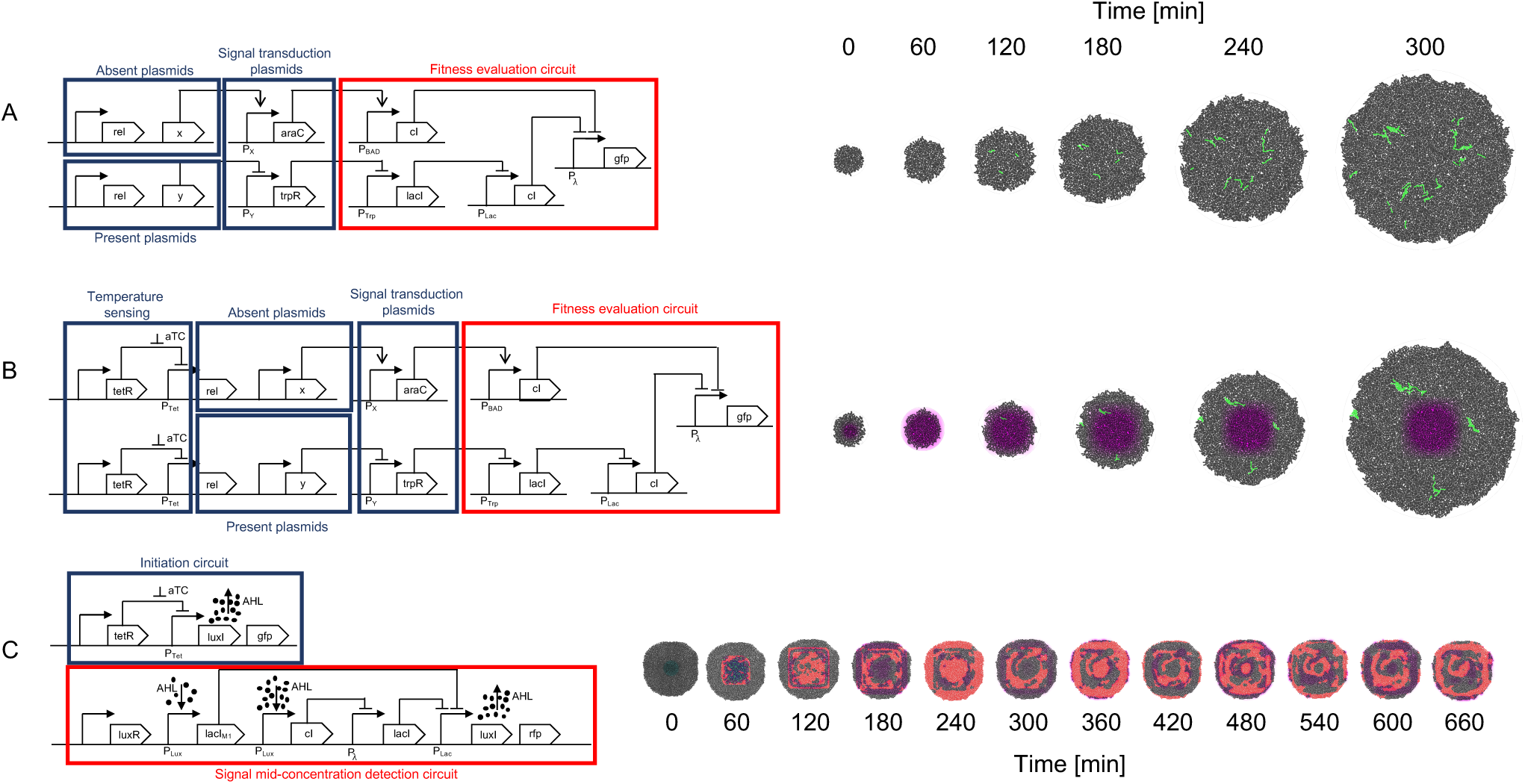
Circuit implementation and simulation. **A**. SGA circuit implementation and simulation. The set of plasmids required to be absent for the fitness function to be optimal are organized in an OR gate manner. If any protein of this set is present, it triggers the cI repressor which in turn represses the fitness reporting (GFP for this example). The plasmids required to be present are processed through an AND gate design: if any of these proteins are missing, the trpR repressor causes the final cI repressor to act on the GFP reporting. Proteins not included in this design are omitted from the fitness evaluation, which is equivalent to a “don’t care” classification in terms of fitness values. All plasmids encoding proteins for evaluation are conjugative. **B**. The SA implementation follows a very similar design to the one made for SGA, however, it incorporates a temperature sensing module, in which aTc concentration regulates conjugation frequency, having higher conjugation rates when aTc concentration is higher, and gradually lower ones as aTc concentration diminishes. **C**. A Game of Life (CA) design is implemented in two parts. It should be noted that a CA does not use a fitness function, therefore does not include this part in the implementation. One of the modules initiates the system, since no AHL signals are present in the beginning (and represent the neighborhood signaling for a “live cell”). Once the system is started, the first module ceases its operation and the other module, an adaptation of the Basu et al.^38^ band detector, evaluates the Game of Life in a mid-range concentration of AHL (for assessing live cells maintaining their state or dead cells coming to life). This second module is the one that keeps the CA running afterwards.

## DISCUSSION

We presented a new framework that proposes a mapping to associate MH to genetic circuits to be encoded in a growing cell colony. The idea behind this proposal is to bring the inspiration from evolution, used for EA (and more broadly MH), back to its origins – an evolving and growing cell colony – for assessing its viability as an implementation testbed. One of the advantages of this approach is that several intrinsic processes involved in the cell and necessary for evolution, such as growth, gene circuit operation, or intercell communication need not be artificially imposed on the model. However, it is their integration into the model, and the level of control which must be studied and adapted to be a suitable element within the mapping. One example of such features may be mutations that occur in the DNA sequence: although this is a process that can be directly linked to mutations in the definition of SGA, for instance, it is also practically impossible to guarantee a mutation rate (as a parameter for SGA). This leads to a couple of possible options. First, a redefinition of MH within a different paradigm immersed in a biological context. Adaptation of MH to a context in which some parameters and/or elements are not fixed, controllable or can be mapped. Second, a direct and artificial mapping of the MH, forcing relationships and mappings to maintain a strict link to their original definition. Since our testbed was a simulation platform, we took a hybrid approach, leaving some of the processes, such as colony growth, to be controlled by the simulator and to be interpreted within the MH execution. This flexibility can be observed, for example, in the implementation of the original version of SA, where there is no mention of “growth” in its definition. Our implementation maps it to the evolution of the temperature function, and thanks to mechanical shoving of the cells outwards of the colony, acts as temperature decrease (aTc concentration is a candidate to represent the temperature measure). However, a strict link was maintained to the evolution of the solution itself and encoded in the cells as a boolean function based on plasmid and/or protein presence. This could have been modeled in a different manner and only have relied on intrinsic mutation. In sum, we propose a framework and one possible mapping for relating MH to synthetic circuits, however, other possible mappings are also valid. Concerning CA, QS is a key player in our implementation, driving the simulation of the model, as it relies heavily on the signal diffusion and degradation parameters. It is very difficult to faithfully reproduce an original Moore neighborhood using QS, since it is not guaranteed that the neighborhood includes a specific number of neighbors or that diffusion can be parametrized, in-vitro or in-vivo, to an extent that an immediate neighborhood can be detected with a very low concentration of AHL. However, we have shown that it is possible to simulate a CA using (simulated) cell colonies. Characterizing the power of CA in cell colonies and specifying the limits of the expressivity for these models becomes an important matter, as there are cases of CA that are Turing-complete models^40,41^.

We believe that the model proposed in this work can be directed towards solving problems involving a large number of variables and in which many solutions need to be evaluated. This is based on the fact that cell colonies have very large counts and large-scale parallelism in the solution evaluations is possible. We think that the number of solutions that is evaluated in parallel within this context is not something that can be achieved by a traditional computer. However, it is also true that even though the proposed model is extensible, a lack of well-characterized synthetic parts may pose a problem in terms of orthogonal intercell communication^42–45^ and variety of elements to construct large synthetic circuits. Future in-vivo implementations can be assisted by software to help select the proper parts for the design^46,47^. Typical SGA and SA algorithms running in a computer use a relatively “low” number of solutions with respect to our proposal (and our testbed is still low on cell counts). Cell colonies are capable of evaluating orders of magnitude more solutions. One constraint to which our work is subjected at this stage is that the current version of the `gro` simulator works with digital proteins. We are aware that this is a large limitation and that the power of the proposed framework would increase greatly by extending its definition to work with analog values of protein expression instead of digital values.

An existing and common related technique is directed evolution. We believe that including rules defined by a MH, evolution can be controlled further and more precisely. Also, it is our opinion that the process is made autonomous, since the selection machinery can be programmed and expressed in terms of synthetic circuits. Also, if multicellular distributed circuits with intercell communication are taken into account, complex computation and conditions can be described^48,49^. Protein engineering^50^ is an application to which this research could be applied in that the properties or functionalities of the protein are encoded as a fitness function, establishing the selection mechanism for desired proteins to be evolved.

### Future work

Current research is being invested into relating different AI algorithms such as Neural Networks, Reinforced Learning (Q-Learning) and other MH, such as Ant Colony Optimization, to our framework. Also, a related direction would be to further study the proposed framework by broadening the tools used to implement the underlying synthetic circuits. An idea in this direction would be to include bacteriophage infection as an intercell communication method^51^ in the framework definition. The current renewed interest in AI, and its need of powerful computational resources, offers a huge opportunity for directing the potential of Synthetic Biology towards satisfying those needs and providing an alternative paradigm (and more natural, since inspiration for most MH actually comes from biology) for solving difficult problems. The goal of this ongoing and future research is to reach the definition of a global AI framework^52^. Characterizing CA computational power in cell colonies is another interesting and important topic to study. Another long-standing debt of our research group is the linkage of the `gro` simulator to accept SBOL^53,54^ specifications as input. In the context of the work presented in this paper, the association of SBOL to Agent/Individual based Model (AbM/IbM) simulators such as `gro` can go further and entail an AI toolkit within SBOL for immediate implementation of such algorithms in cell colonies. A more distant direction is to use the framework to internally implement EA for Xenobots^55^ using division of labor^56^ to perfect and automate their functionality.

### Materials and Methods

We used a new version of `gro` developed by AI-UDP for the simulations. This new version can be found at https://github.com/AI-UDP/GRO63. All simulations (`gro`, C, and C++) were run in MacOS Catalina version 10.15.2 and in Windows 10. The interpreter to generate the `gro` simulation files was written in C++ and can be found at https://github.com/AI-UDP/MHInterpreter. Machines used for simulations were two MacBook Pro core i5 2.7 GHz and 2.5 GHz with 8GB RAM and a Pentium G4560 3.GHz with 16GB RAM.

## ASSOCIATED CONTENT

### Supporting Information

The following files are available free of charge.

**Text S1**. Data for both implementations of SA and SGA, and Conway’s Game of Life. More information on the interpreter is also provided.

**SA.mp4**. Video of the Simulated Annealing execution in `gro`.

**SGA.mp4**. Video of the Simple Genetic Algorithm execution in `gro`.

**CA.mp4**. Video of the Cellular Automaton in `gro`.

**SA.gro** – This is the file generated (.gro) by the interpreter and used in the `gro` simulator for Simulated Annealing (SA).

**SGA.gro** – This is the file generated (.gro) by the interpreter and used in the `gro` simulator for Simple Genetic Algorithms (SGA).

**CA.gro** – This is a file we implemented (.gro) and used in the `gro` simulator for Conway’s Game of Life.

**SA.c** – This is a source file implemented in C to generate data and compare with SA.gro data.

**SGA.cpp** – This is a source file implemented in C++ to generate data and compare with SGA.gro data.

**gameOfLife.c** – C implementation of a cellular automaton (Game of Life) to compare its execution with CA.gro.

## Author Contributions

Framework design: MGP, Interpreter design and implementation: JCR, Design and implementation of the `gro` versions of the algorithms: MGP, JCR, Design and implementation of C and C++ versions of the algorithms: YOM, Synthetic circuit designs: MGP, JCR, YOM, Simulation executions and data processing: JCR, YOM, Wrote the paper: MGP, Supplementary information document: YOM, MGP

The manuscript was written through contributions of all authors. All authors have given approval to the final version of the manuscript.

## ACKNOWLEDGMENT

The authors thank Luciano Ahumada, Marco Clavero, Sebastián Antón and Pablo Ramos for their comments and valuable discussions, Guillermo Iglesias for helping in automating and running the SA `gro` simulations, Aaron Adler and Fusun Yaman for their valuable insight in the initial stages of this work at the AI for Synthetic Biology Workshop, 2018. The authors are also grateful for all feedback received at IWBDA 2019 for developing this work.

