## Supplemental data for "A framework for implementing metaheuristic algorithms using intercellular communication"

### Supporting Information

#### Simulation data for original and framework (`gro`) versions of MH

##### Simple genetic algorithm (SGA)

One version of the SGA was implemented in C++17 and seeks to create a population of individuals of size N, in such a way that it finds the individuals that possess a given distribution of K plasmids. The presence of every plasmid is encoded in a bit string: if the $i^{th}$ plasmid is present in the individual then the $i^{th}$ bit is a 1, for the complementary case the $i^{th}$ bit is a 0. Another implementation was generated for the `gro` simulator. Under this implementation, mutation represented as an abnormal function of the promoter, crossover is implemented as bacterial conjugation and the fitness function is checked by evaluating the presence, absence or indifference of each plasmid in a set within a bacterium. We compared number of generations until finding the first instance of an optimal solution, because other measures are difficult to associate given differences in operation implementations and parametrization.

The parameters of the program are the population size, mutation rate, crossover rate, number of plasmids, the fitness function (in the form of a bit sequence that represents the plasmids of the optimal individual), and the maximum number of iterations of the algorithm. It is important to highlight that the population in this program is static, it does not grow as in the `gro` simulation. Tests were performed on the C++ simulations by varying the fitness function to evaluate, the mutation rate, crossover rate, and population size. The corresponding simulations were run in the `gro` implementation. The baseline configuration for all simulations was: 1100 fitness function, 1% crossover rate, 1% mutation rate and 200 initial solutions. Using this baseline configuration, respective parameters were varied to get the results for the execution of SGA. The initial populations for all simulations were composed by 50% of empty solutions (holding no plasmids initially) and 50% holding a single plasmid, distributed equally.

Tests were carried out for four different fitness functions. For each fitness function, 30 simulations were carried out.

| Fitness function | First appearance of optimal solution (generation nº) – `gro` | First appearance of optimal solution (generation nº) – C++ |
|---|---|---|
| 11111 | Solution not found | 7.8 |
| 11000 | 2.293 | 3.840 |
| 1100 | 2.199 | 3.963 |
| 10xxx | 0.396 | 0 |

**Table 1.** Data of the mean number of generations in which the first optimal solution appears in the C++ versions of the simulation. The fitness function is expressed as a 1 when the plasmid is required to be present, while a 0 requires the plasmid to be absent. An x represents indifference for whether the plasmid is present or absent.

The results in Table 1 show that finding an optimal solution takes longer when a more restrictive function is sought. In the context of this problem, the term restrictive means that more plasmids must be present simultaneously in the solution. It should be noted that 10xxx is practically found upon algorithm execution. This is because the solution 10000 is part of the initial solution pool, and it matches the fitness function 10xxx. In the case of the `gro` implementation, its detection is not immediate, because it has to express the protein signaling the plasmid presence.

Then, mutation rate was varied in the executions. 30 simulations were also run for each of the mutation rate settings.

| Mutation rate | First appearance of optimal solution (generation nº) – `gro` | First appearance of optimal solution (generation nº) – C++ |
|---|---|---|
| 0% | 3.303 | Solution not found |
| 2.5% | 3.194 | 4.5 |
| 5% | 2.060 | 1.5 |
| 10% | 1.27 | 1.0 |

**Table 2.** Mean number of generations it takes to find the first optimal solution by varying the mutation rate.

Mutation rate configures how often a mutation occurs. In SGA, mutations represent global search operations. Therefore, a higher mutation rate speeds up finding optimal solutions, but it can affect local search sequence. However, it should be noted that a mutation rate of 0 is unable to find an optimal solution.

30 simulations were also carried out for each crossover rate value.

| Crossover rate | First appearance of optimal solution (generation nº) – `gro` | First appearance of optimal solution (generation nº) – C++ |
|---|---|---|
| 0% | 3.745 | 3.821 |
| 2.5% | 3.328 | 3.286 |
| 5% | 1.978 | 4.679 |
| 10% | 1.27 | 2.727 |

**Table 3.** Mean number of generations it takes to find the first optimal solution by varying the crossover rate in both SGA implementations.

Table 3 shows that the number of generations required to find the optimal solution tends to decrease as the crossover rate increases. Unlike mutation, the crossover operation implements local search in the region of the search space in which the original solutions involved in recombination are located.

Finally, 10 executions of each initial population size were carried out.

| Population size | First appearance of optimal solution (generation nº) – `gro` | First appearance of optimal solution (generation nº) – C++ |
|---|---|---|
| 200 | 3.187 | 4.011 |
| 400 | 2.162 | 3.942 |
| 10000 | 0.596 | 1.463 |

**Table 4.** Data for the mean number of generations before finding an optimal solution by varying the size of the initial population of solutions.

The results for population size in Table 4 show that if the initial population is larger, then less generations are needed to find an optimal individual.

Although numbers for the generated data are different (this is due to differences in representation and mechanics involved in specific operations of the algorithm), the behaviour shown by both kinds of simulations (`gro` and C++) is comparable. Changes in the parameters exhibit similar output result changes.

**Simulated Annealing (SA)**

Like in SGA, both the C and the `gro` version of SA perform optimization to find an individual holding a given distribution of K plasmids. In this model, the presence of a plasmid is encoded as a 1 at the respective location in the bit sequence representing the whole set of plasmids. Conversely, whenever the absence of the plasmid is required, it is expressed as a 0 at the location corresponding to the plasmid in question. It is also possible to be indifferent to the presence of a given plasmid, in which case the plasmid is not mentioned in the distribution.

The C version of the SA algorithm receives the following parameters: fitness function (expressed in a similar fashion than in SGA – sets of present, absent and indifferent plasmids), number of plasmids, initial temperature, the minimum temperature that marks the end of the execution of the program, and alpha (a value that denotes the decrease rate in temperature). For this instance of the model, the temperature decrease is encoded as a linear function with the alpha value being strictly lower than one. It should be noted that this version of SA works with a single solution. In contrast, the `gro` version of SA uses several solutions (each one is an individual bacterium). Therefore, an additional parameter is required: initial population size. The fitness function under this implementation is expressed a set of plasmids that need to be present, absent or their presence is indifferent. Finally, the temperature decrease (alpha) is translated to a degradation rate of the environmental signal. A baseline configuration for simulation was also set for SA: a value of 0.25 for alpha and a fitness function of 1100. For the `gro` version, an additional default value of 200 was set for the initial population. As in SGA, initial populations for all simulations were made up of 50% of empty solutions (holding no plasmids initially) and 50% holding a single plasmid, distributed equally for the `gro` version.

Our team ran tests varying the fitness function and temperature decrease rate. First, we varied the fitness function, executing 200 times each simulation with a specific configuration for the C version and 30 times for the `gro` version.

| Fitness function | First appearance of optimal solution (generation nº) – `gro` | First appearance of optimal solution (generation nº) – C |
|---|---|---|
| 11111 | Solution not found | 8.3 |
| 11000 | 3.238 | 8.45 |
| 1100 | 5.626 | 8.83 |
| 10xxx | 0.7499 | 10.04 |

**Table 5.** Mean number of generations for each configuration to find the first instance of an optimal solution. This is done for different fitness functions.

Table 5 shows stable values for the required number of generations to find the optimal solution using the C version of the algorithm. This can be explained by the fact that the initial solution is random, and the value change of a 0 to a 1 depends on the temperature value. However, the `gro` version of the algorithm behaves similar to what was observed in SGA. This is, the more restrictive a fitness function is, the longer it takes to find an optimal solution.

Other simulations were run for different configurations of temperature decrease rates (alpha). As in the case of the fitness function, 200 repeats were performed for each alpha configuration (C version) and 30 repeats for each degradation rate (`gro` version).

| Degradation rate / alpha | First appearance of optimal solution (generation nº) – `gro` | First appearance of optimal solution (generation nº) – C++ |
|---|---|---|
| 0.25 | 3.510 | 8.83 |
| 0.5 | 2.321 | 21.83 |
| 0.75 | 5.004 | 22.77 |
| 0.9 | 3.209 | 22.76 |

**Table 6.** Mean number of generations required to find the first instance of an optimal solution. Several temperature decrease rates were tested.

Results from Table 6 show a large variability in the `gro` version of the algorithm. Our explanation is that achieving an optimal solution depends mostly on the spatial location of the multiple solutions, therefore due to randomness in this location and the influence of the temperature decrease occurring outwards of the colony, no tendency can be accurately found in these cases. However, in the C++ version of the algorithm, an increase in alpha entails an increase in the generations for finding the first optimal solution. This is due to a single solution being explored, and therefore, evolution being slowed down as the temperature decrease rate is greater.

**Game of Life – Cellular automaton (CA)**

Our team implemented a version of Conway's Game of Life in C. It is a grid of n x m cells where every cell has two possible states, live or dead. The state changes are governed by the following rules: if a dead cell has two or three live cells in its Moore neighborhood then this cell will come back to life, but if a cell has more than 3 or less than two live cells in its Moore neighborhood then this cell will die by overpopulation or underpopulation respectively. An adaptation of this model was implemented in `gro` using the same rules.

For this model, the only results that our team compared were the spatial patterns that the automaton exhibits. This comparison was made between the patterns generated by the C version and the `gro` version.

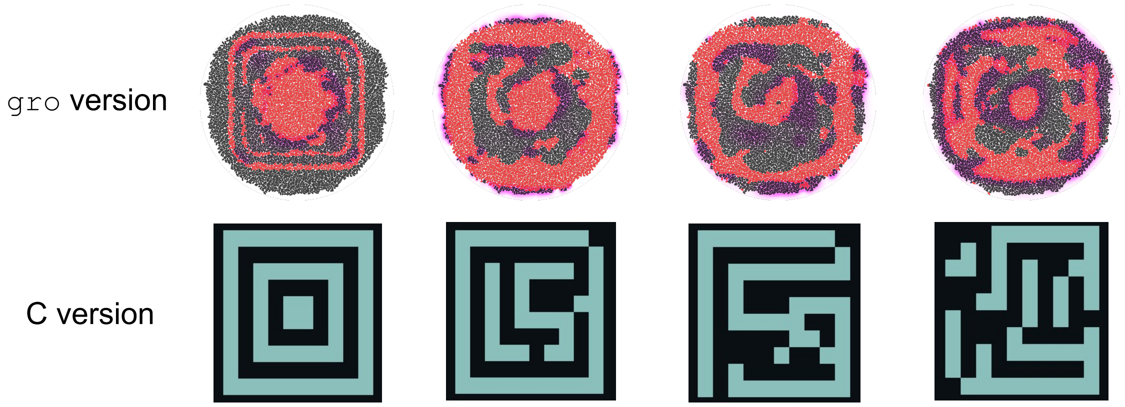


**Figure S1.** Comparison of spatial patterns achieved by Conway's Game of Life implementations in `gro` and in C. In the `gro` implementation, red cells represent live ones, and uncolored cells represent dead ones. The magenta regions are locations at which there is a high concentration of AHL. In the C implementation, cyan cells are live ones and black cells are dead ones.

Although the patterns shown in Figure S1 are not identical, they are similar in their shape. Dimension discrepancies can be attributed to the difference in the calculation of the Moore neighborhood for both versions, as the `gro` version uses AHL signal to find and sense its neighborhood, while the C version of the neighborhood is calculated directly on a grid and does not present variability. Also, the AHL concentration used for defining the neighborhood must be parametrized, and depending on the values of emission and sensing, it can cause the neighborhood to extend past immediate contiguous cell neighbors, turning a cell group into what should be a single grid cell. Finally, it has not been possible for our simulations to achieve perfect fine-grained equivalence between a bacterium and a grid cell, explaining why the patterns exhibit differences.
